# IL-27 Signaling Protects Against Influenza-Associated Pulmonary Aspergillosis Through Inhibition of Type 2 Immunity and Enhanced Antifungal Immunity

**DOI:** 10.64898/2026.07.10.737799

**Authors:** Nima Naghshtabrizi, Yonne Karoline Tenório de Menezes, Shuxia Wang, Ravineel B. Singh, Dequan Lou, Kong Chen, Radha Gopal, Caden Ngeow, Keven M. Robinson

## Abstract

Influenza-associated pulmonary aspergillosis (IAPA) is a severe complication of influenza infection associated with substantial mortality. Influenza disrupts pulmonary host defenses and alters innate immune responses, predisposing patients to invasive fungal infection. Interleukin-27 (IL-27) is an immunoregulatory cytokine with context-dependent antiviral and antifungal effects; however, its role during IAPA remains undefined. A mouse model of IAPA was established by infecting wild-type and IL-27 receptor α-deficient (*Il27ra^−^/^−^*) mice with influenza A, followed by *Aspergillus fumigatus* challenge. IL-27 and IL-27Rα expression were increased during IAPA. Single-cell RNA sequencing identified monocytes as the primary source of IL-27 and T cells as major IL-27rα-expressing cells. *Il27ra^−^/^−^* mice exhibited significantly increased pulmonary fungal and influenza viral burden, enhanced type 2 immune responses characterized by elevated IL-4, IL-5, IL-9, IL-13, eosinophils, Th2 cells, pathogenic Th2 cells, and ILC2s. Despite increased eosinophil abundance, eosinophil-mediated conidial killing was impaired in *Il27ra^−^/^−^* mice. IL-27Rα deficiency also reduced macrophage abundance and impaired macrophage conidial uptake. Conversely, timed administration of rIL-27 enhanced fungal clearance, improved survival, and increased macrophage conidial uptake and augmented eosinophil killing capacity during IAPA. IL-27 signaling is a protective immunoregulatory cytokine during IAPA that limits pathological type 2 inflammation and enhances antifungal effector function of both eosinophils and macrophages. These findings identify IL-27 as a potential therapeutic in IAPA.

## Introduction

Influenza remains a substantial global health burden, causing approximately 1 billion infections annually and an estimated half a million respiratory-related deaths each year(1, 2). In critically ill patients, influenza-associated pulmonary aspergillosis (IAPA) is increasingly recognized as a frequent and highly lethal complication, with reported intensive care unit incidences of ~10–19% and a 90-day mortality of ~50%, substantially exceeding the mortality associated with severe influenza alone(3–5). Influenza infection can predispose to invasive pulmonary aspergillosis by disrupting pulmonary host defenses, including the epithelial cell barrier, mucociliary clearance, immune cell recruitment, and immune cell function(6, 7). We have previously shown that influenza-driven signaling reduces alveolar macrophage abundance and limits neutrophil accumulation in the lung, in part through a signal transducer and activator of transcription 1 (STAT1)-dependent pathway(8). While innate immune defects during IAPA are increasingly being recognized, it remains unclear how immunoregulatory pathways engaged during influenza reshape the inflammatory milieu and influence antifungal immunity during IAPA.

Multiple cytokines and interferons, including IL-27, signal through the transcription factor STAT1(9, 10). IL-27 is a heterodimeric immunoregulatory cytokine that is made up of IL-27p28 and EBV-induced gene 3 (EBI3) and regulated by activation of toll-like receptors (TLRs) and nuclear factor kappa B (NF-kB) signaling pathways(11)(12). The IL-27 receptor (IL-27R) is composed of two subunits, IL-27Rα (*Il27ra*) and gp130 (*Il6st*)(13). IL-27 signals through the IL-27R to activate Jak1/Jak2 and phosphorylation of STAT1/STAT3 transcription factors, mediating downstream effects(14–16). IL-27 promotes Th1 differentiation while restraining type 2 inflammation, including via suppression of innate lymphoid cell (ILC)2 expansion and cytokine production, and reduced eosinophilic airway inflammation(17–19). IL-27 promotes both pro-inflammatory and anti-inflammatory responses, and these responses can be context-dependent. IL-27 has potent antiviral properties; IL-27 upregulates during influenza infection in both mice and humans(20, 21). IL-27 is also upregulated during invasive pulmonary aspergillosis and impairs host defense against *Aspergillus fumigatus* (*A. fumigatus*) with decreased fungal burden and improved survival(22). The role of IL-27 signaling during IAPA remains undefined, and we herein utilize a IAPA mouse model to profile immune responses during IAPA with the goal of identification of new therapeutics.

## Materials and Methods

### Animal Models

Male C57BL/6 wild-type mice (6–8 weeks old) were purchased from Taconic Farms (Germantown, NY). IL-27 receptor α-deficient (*Il27ra^−/−^*) mice on the C57BL/6N background (B6N.129P2-Il27ra^tm1Mak^/J) were purchased from Jackson Laboratories and bred at the University of Pittsburgh for experimental use. All mice were housed under specific pathogen-free conditions at the University of Pittsburgh animal facilities and acclimatized prior to experimentation. Studies were conducted using age- and sex-matched male mice in accordance with protocols approved by the University of Pittsburgh Institutional Animal Care and Use Committee.

### Pathogens

Mice were inoculated with influenza A/PR/8/34 (H1N1), generously provided by Dr. Radha Gopal (UPMC Children’s Hospital of Pittsburgh), at a sublethal dose of 2000 plaque-forming units (PFU) in 50 µL sterile PBS via oropharyngeal aspiration. This inoculum consistently produced 15–20% body weight loss by day 6 post-infection, serving as a model of severe influenza. Pulmonary viral burden was quantified by RT-qPCR targeting the viral matrix protein on lung-derived RNA. *A. fumigatus* (ATCC 42202) was cultured on potato dextrose agar at 37°C for 5–7 days. Conidia were harvested by washing culture flasks with 25 mL of sterile PBS containing 0.1% Tween 20 and quantified using a hemocytometer. For fungal burden quantification, the right upper lobe of each lung was mechanically homogenized in 1 mL sterile PBS. Homogenates were diluted 1:1000 and 100 µL was plated on potato dextrose agar for colony-forming unit (CFU) enumeration. The <u>fl</u>uorescent *<u>A</u>spergillus* <u>re</u>porter (FLARE) strain (Af293-dsRed) was kindly provided by Dr. Tobias Hohl (Memorial Sloan Kettering Cancer Center). FLARE conidia were prepared as previously described(23). 1 × 10^8^ Af293-dsRed conidia were incubated with 0.5 mg/mL Biotin-XX, SSE (ThermoFisher) in 50 mM carbonate buffer (pH 8.3) at 4°C for 2 hours on a tube rotator, washed with 0.1 M Tris-HCl, and subsequently labeled with 0.02 mg/mL streptavidin-Alexa Fluor 647 (AF-647, Life Technologies) for 30 minutes at room temperature protected from light with rotation. Labeled conidia were resuspended in PBS with 0.025% Tween 20.

### Bronchoalveolar Lavage

Bronchoalveolar lavage (BAL) was performed by instilling and recovering 1 mL of sterile PBS through the trachea. BAL fluid was centrifuged for 5 minutes at 4°C to pellet cells. The cell pellet was resuspended and total cell counts were determined using a hemocytometer. Cyto-spin preparations were made from BAL cells and stained for differential cell counting to quantify immune cell populations. Total protein concentration in cell-free BAL supernatant was measured as a surrogate marker of alveolar-capillary barrier injury using the Pierce BCA Protein Assay Kit (Thermo Scientific, catalog #23225) according to the manufacturer’s instructions. Briefly, 50 µL of BAL fluid was added per well in duplicate alongside a bovine serum albumin standard curve prepared by twofold serial dilution. BCA working reagent was added to each well and incubated for 15 minutes at 37°C. Absorbance was measured at 562 nm on a plate reader and protein concentration calculated using a four-parameter logistic standard curve.

### Multiplex Cytokine Analysis

The cranial lobe of the right lung was homogenized in sterile PBS by mechanical grinding. Simultaneous quantification of multiple cytokines and chemokines was performed on lung homogenate using the Bio-Plex Pro Mouse Cytokine 23-plex Panel (Bio-Rad Laboratories, #M60009RDPD) on the Bio-Plex 200 system per manufacturer’s instructions, with data analyzed using a five-parameter logistic standard curve.

### RNA Extraction and RT-qPCR

The middle and lower lobes of the right lung were rapidly frozen and homogenized in liquid nitrogen prior to RNA extraction using an RNA isolation kit (Agilent Technologies). RNA expression levels were subsequently analyzed using standard RT-qPCR with Bio-Rad SSO Advanced Universal Probes Supermix (CA, USA). Gene expression was assessed in technical duplicate. Relative expression levels were calculated using the formula ΔCq=2^Cq^ ^target^ ^gene-Cq^ ^ref^, where *Cq* represents the mean quantification cycle value of the target gene normalized to the average value of the HPRT reference gene.

### Histology

Whole-slide GMS images produced from the whole left lung and were imported into QuPath and analyzed using a supervised pixel-classification workflow to identify fungal conidia as black/silver-stained punctate structures against the counterstained lung background(24). Representative regions of interest were manually annotated to capture staining variability and artifacts, and pixels were labeled into classes of conidia vs background lung tissue. A QuPath pixel classifier was trained with live prediction enabled, and training regions were iteratively expanded until the overlay consistently detected conidia while minimizing false positives from nonspecific dark elements. The finalized classifier was saved and applied uniformly across all sections to generate a conidia mask; minor post-processing was used as needed to remove spurious detections, and conidial burden was reported as the frequency of conidia normalized to tissue to enable comparisons across samples.

### Survival curve

Body weight was monitored twice daily. Mice that lost ≥25% of their initial body weight were humanely euthanized per University of Pittsburgh IACUC guidelines, and this time point was recorded as the survival endpoint. Kaplan-Meier survival curves were generated, and statistical differences between groups were determined by the log-rank (Mantel-Cox) test.

### Flow Cytometry

To create single-cell suspensions from lung tissue, enzymatic digestion was performed using collagenase IV and DNase, followed by mechanical separation through a 70-µm cell strainer. Red blood cells were eliminated using ACK lysis buffer. The cells were then counted and resuspended in PBS. To prevent non-specific antibody binding, anti-mouse TruStain FcX™ CD16/CD32 (Biolegend) was applied for 10 minutes at 4°C. Surface staining was conducted with fluorochrome-conjugated antibodies for 30 minutes at 4°C, shielded from light. Cell viability was evaluated using Live/Dead Fixable Blue or Zombie NIR viability dye according to the manufacturer’s guidelines. After surface staining, cells were fixed with paraformaldehyde. For intracellular targets, cells underwent permeabilization and staining following standard protocols. Data acquisition was carried out on a Cytek Aurora spectral flow cytometer, and analysis was performed using FlowJo software (BD Biosciences). Gating strategies for all cell populations, including FLARE-based antifungal functional assays and antibodies are provided in supplemental data (Figure S1, Table S1). Absolute cell counts were calculated by normalizing flow cytometry event counts for each population to the total cell count determined by automated counting prior to staining, adjusted for the fraction of cells loaded per well.

### RNA Sequencing

#### Single-Cell RNA Sequencing

Single-cell RNA sequencing data were generated from CD45^+^ immune cells isolated from mouse lungs using CD45 MicroBeads (Miltenyi Biotec). Reads were aligned to the mouse reference genome (mm10/GRCm38) and quantified using Cell Ranger (v7.0; 10x Genomics). Downstream processing and cell clustering were performed in Seurat (v5.2.0), with cell identities assigned based on canonical lineage markers across myeloid and lymphoid compartments. Expression of *Il27*, *Ebi3*, *Il27ra*, and *Il6st* was examined across identified immune cell populations.

#### Bulk RNA Sequencing — Whole Lung Homogenate

For transcriptomic profiling, RNA was extracted from snap-frozen whole lung homogenates using the RNA isolation Kit (Agilent Technologies) per the manufacturer’s protocol. RNA quantity and integrity were assessed prior to library preparation. Library preparation was performed and sequencing was carried out the Health Sciences Sequencing Core at UPMC Children’s Hospital of Pittsburgh. For the comparison of WT Flu/AF versus AF-only mice, pathway analysis was performed using QIAGEN Ingenuity Pathway Analysis (IPA, version 127006219). Pathway activation Z-scores were calculated from expression log ratios of differentially expressed genes, where positive values indicate pathway activation in the first group relative to the second. For the comparison of IL-27Rα^−^/^−^ Flu/AF versus WT Flu/AF mice, pathway analysis was performed using IPA as described above. GSEA was additionally performed using the MSigDB Hallmark gene sets (*Mus musculus*) via the R package msigdbr, with genes ranked by average log₂ fold-change.

#### Recombinant IL-27 Administration

To assess the effect of IL-27 augmentation during IAPA, wild-type C57BL/6 mice received intraperitoneal injections of recombinant murine IL-27 (rIL-27; 200 ng in 100 µL sterile PBS; BioLegend, catalog #577404) on days 4–7 of the IAPA model. Control mice received an equivalent volume of sterile PBS by intraperitoneal injection on the same days. Mice were challenged with *A. fumigatus* on day 6 post-influenza infection and lungs were harvested 48 hours post-fungal challenge.

## Results

### IL-27 signaling is increased during IAPA

We have previously shown that preceding influenza A infection increases susceptibility to secondary *A. fumigatus* super-infection in the mouse lung and allows for the development of IAPA with increased fungal burden, increased fungal germination, cellular inflammation, lung injury, and mortality(8). To further investigate how influenza alters antifungal host defense, C57BL/6 male mice were challenged with a sublethal dose of influenza A PR/8/34 H1N1 (2000 pfu) for 6 days, followed by 2.5 × 10^7^ *A. fumigatus* (ATCC strain 42202) conidia, and after 48 hours, lungs were harvested and RNA sequencing performed. Mice were challenged with *A. fumigatus* on day 6 post-influenza infection to model IAPA that occurs in critically ill patients hospitalized with severe influenza(25). The greatest difference in fungal burden between IAPA and singular *A. fumigatus* infection in our model occurs at 48 hours post-fungal challenge(8), and therefore, we pursued lung RNA sequencing at this time point. RNA sequencing pathway analysis demonstrated an increase in Type I and Type II interferon, IL-27, Type 1 and Type 2 immune signaling, and IL-10 pathways in IAPA compared to mice infected with *A. fumigatus* alone (Figure 1A). We have previously shown that influenza alters innate antifungal host defense, in part through a STAT1-dependent pathway. Several influenza-induced interferons and cytokines signal through STAT1, including IL-27. We observed increased expression of *Il27* and *Il27rα* in IAPA compared to *A. fumigatus* infection alone (Figure 1B). To determine the kinetics of IL-27 expression during IAPA, and validate our RNA sequencing findings, we challenged mice with influenza A followed by *A. fumigatus* and harvested lungs at 18 – 96 hours post-fungal challenge and observed a persistent increase in IL-27 expression in IAPA mice compared to *A. fumigatus* alone (Figure 1C). To determine the cellular source of IL-27 and IL-27R subunits in immune cells, we isolated CD45^+^ cells from the lungs of IAPA and singular *A. fumigatus* infected mice and performed single cell RNA sequencing. *Il27* was predominantly expressed in monocytes and *Il27ra* is predominantly expressed in T cells. There is higher expression of *Il27* in IAPA monocytes and higher expression of *Il27ra* in IAPA T cells compared to singular *A. fumigatus* (Figure 1D). Notably, there are non-immune cells that will express the components of IL-27 and IL-27R that were not included in this analysis. To confirm our transcriptomic findings at the protein level, we measured IL-27p28 using flow cytometry and observed increased numbers of IL-27p28^+^ monocytes, neutrophils, alveolar macrophages, interstitial macrophages, and dendritic cells in IAPA compared to singular *A. fumigatus* (Figure 1E). These data demonstrate that IL-27 and the IL-27R are increased during IAPA in both mice and humans. Monocyte-derived IL-27 is upregulated during IAPA, with T cells appearing as the major target of IL-27 signaling.

**Figure 1.**
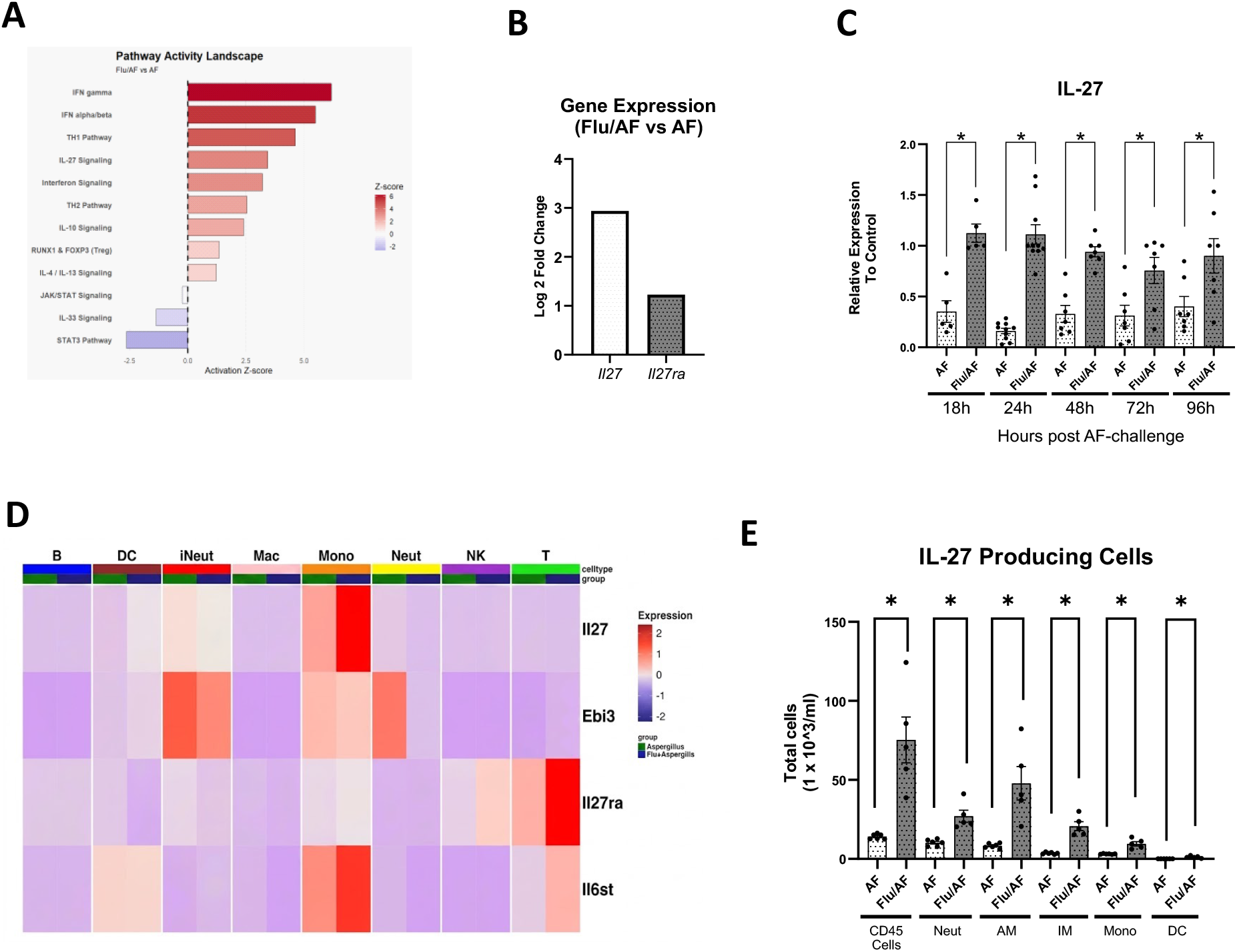
IL-27 signaling is increased during IAPA in mice and humans. A-Pathway activity landscape from bulk RNA sequencing of whole lung homogenates comparingFlu/AF (IAPA) to AF-only infected mice at 48 hours post-fungal challenge. Activation Z-scores are shown for selected immune signaling pathways; red indicates pathway activation and blue indicates inhibition (n= 4 mice per group, data from one experiment).B-Log2 fold change in *Il27* and *Il27ra* gene expression in IAPA compared to AF-only infected mice by bulk RNA sequencing of whole lung homogenates at 48 hours post-fungal challenge (n= 4 mice per group, data from one experiment).C-Relative *Il27* mRNA expression in lungs of AF-only and Flu/AF (IAPA) mice at 18, 24, 48, 72, and 96 hours post-AF challenge, measured by qPCR and normalized to control (n= 5-10 mice per group).D-Heatmap of *Il27*, *Ebi3*, *Il27ra*, and *Il6st* expression across immune cell populations identified by single-cell RNA sequencing of CD45^+^ lung cells from AF-only (Aspergillus) and Flu/AF (IAPA) mice. Cell types shown: B cells, dendritic cells (DC), inflammatory neutrophils (iNeut), macrophages (Mac), monocytes (Mono), neutrophils (Neut), NK cells, and T cells (n= 2 mice per group, data from one experiment).E-Total numbers of IL-27p28^+^ cells among CD45^+^ cells, neutrophils, alveolar macrophages, monocytes, and dendritic cells in lungs of AF-only and Flu/AF (IAPA) mice quantified by flow cytometry (n= 5 mice per group, data from one experiment). Data are shown as means ± SEM. *p<0.05, **p<0.005, ***p<0.0005, ****p<0.0001 by two-way ANOVA with Tukey’s multiple comparisons test (C) and unpaired t-test (D). All experiments represent a minimum of two independent experiments unless otherwise noted.

### IL-27Rα deficiency increases pathogen burden during IAPA

To assess the impact of IL-27 signaling during IAPA, we challenged wild-type (WT) C57BL/6 and *Il27ra^−/−^* mice with influenza A PR/8/34 H1N1 for 6 days followed by *A. fumigatus* conidia and after 48 hours, measured fungal burden in the lung. We observed increased fungal burden in the IAPA *Il27ra^−/−^* mice compared to IAPA WT controls (Figure 2 A) through measurement of colony-forming units (CFU) in lung homogenate. Histologic analysis using Grocott-Gomori methenamine silver (GMS) staining of lung tissue also revealed increased *A. fumigatus* fungal burden in IAPA *Il27ra^−/−^* mice compared to IAPA WT controls (Figure 2 B-C). Additionally, we observed increased influenza viral burden in the lungs of IAPA *Il27ra^−/−^* mice (Figure 2 D). We also observed increased total protein levels and increased numbers of neutrophils in the airway compartment in IAPA *Il27ra^−/−^* mice compared to IAPA WT controls (Figure 2 E-F).

**Figure 2.**
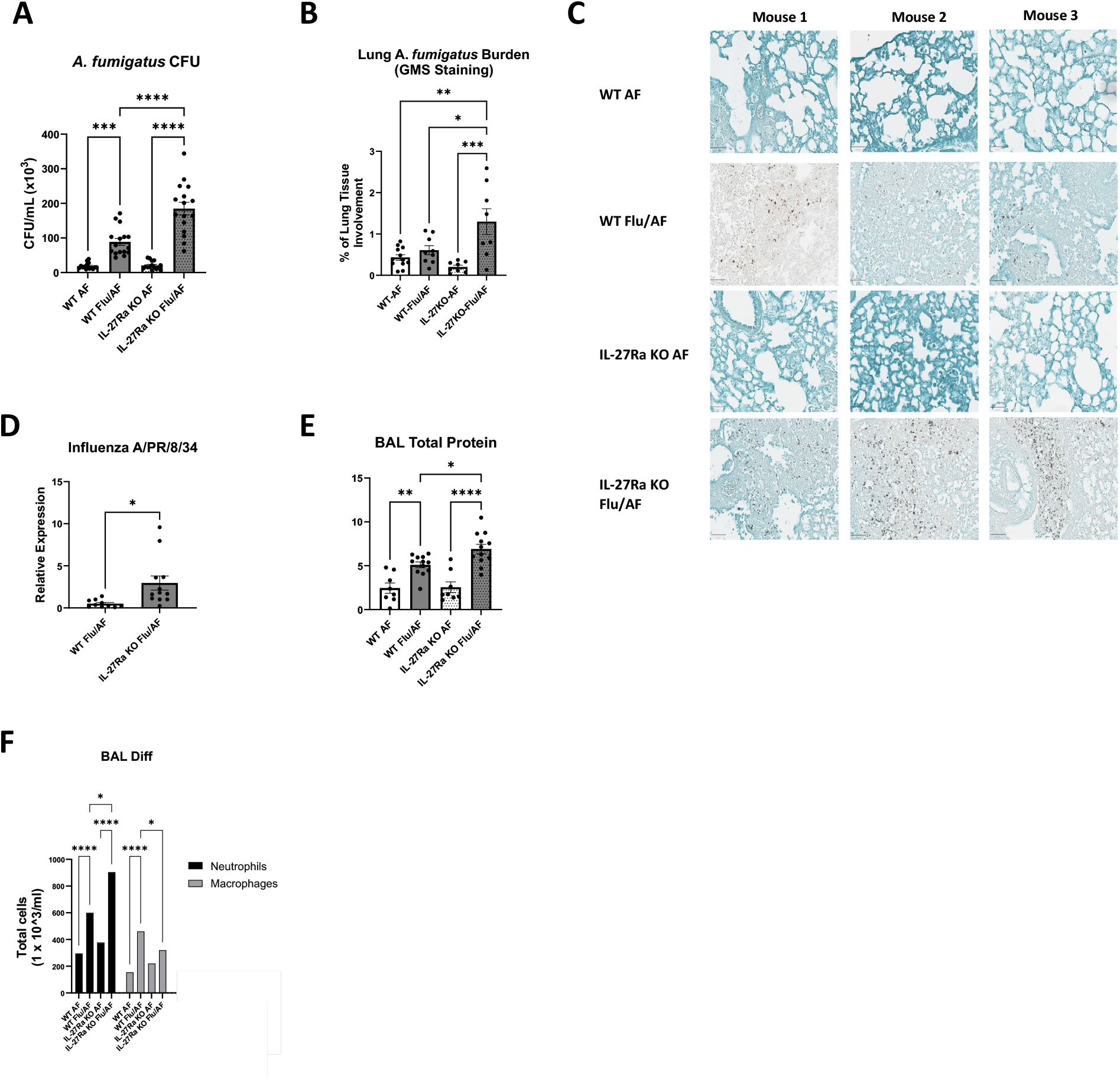
IL-27Rα deficiency increases pathogen burden during IAPA. A-*A. fumigatus* colony-forming units (CFU/mL ×10^3^) in lung homogenates of WT AF, WT Flu/AF, IL-27Rα KO AF, and IL-27Rα KO Flu/AF mice at 48 hours post-AF challenge (n= 16 mice per group).B-Digital quantification of pulmonary *A. fumigatus* burden on Grocott-Gomori methenamine silver (GMS)-stained lung sections, expressed as percentage of lung tissue area occupied by fungal elements, across the four groups (n = 8–12 mice per group). C-Representative GMS-stained lung sections from three mice per group. D-Relative influenza A/PR/8/34 viral burden by qPCR in lungs at 48 hours post-AF challenge. E-Total protein concentration in bronchoalveolar lavage fluid (n = 8–12 mice per group). F-Total numbers of neutrophils and macrophages in bronchoalveolar lavage fluid quantified by cytospin differential cell counting (n= 8 mice per group). Data are shown as means ± SEM. *p<0.05, **p<0.005, ***p<0.0005, ****p<0.0001 by one-way ANOVA with Tukey’s multiple comparisons test (A–B) and unpaired t-test (D). All experiments represent a minimum of two independent experiments unless otherwise noted.

### IL-27Rα deficiency enhances Type 2 Immunity but limits eosinophil-mediated conidial killing during IAPA

To examine the host response during IAPA, we performed bulk RNA sequencing on the lung tissue of IAPA *Il27ra^−/−^* mice compared to WT. Pathway analysis demonstrated upregulation of the TH2, IL-4/IL-13 signaling, IFN gamma, and IFN alpha/beta pathways and downregulation of the IL-10 signaling pathway in IAPA *Il27ra^−/−^* mice compared to IAPA WT controls (Figure 3 A). We measured protein levels of cytokines, chemokines, and interferons in the lung during IAPA in WT and *Il27ra^−/−^* mice. We observed increased IL-4, IL-5, IL-13, IL-9, eotaxin, and GM-CSF in the IAPA *Il27ra^−/−^* mice compared to IAPA WT controls (Figure 3 B-G). Interestingly, IL-33 gene expression was unchanged in IAPA *Il27ra^−/−^*mice compared to IAPA WT controls; however, it was highly upregulated during *A. fumigatus* infection in *Il27ra^−/−^* mice compared to WT controls (Figure 3H). We observed increased levels of IFN-γ in IAPA *Il27ra^−/−^* mice compared to IAPA WT controls (Figure 3I). IL-17 levels were significantly higher in IAPA *Il27ra^−/−^* mice compared to IAPA WT controls, despite IFN-γ levels also being elevated in the *Il27ra^−/−^* mice (Figure 3J). IL-10 levels were unchanged between the WT and *Il27ra^−/−^* mice IAPA mice (Figure 3K).

**Figure 3.**
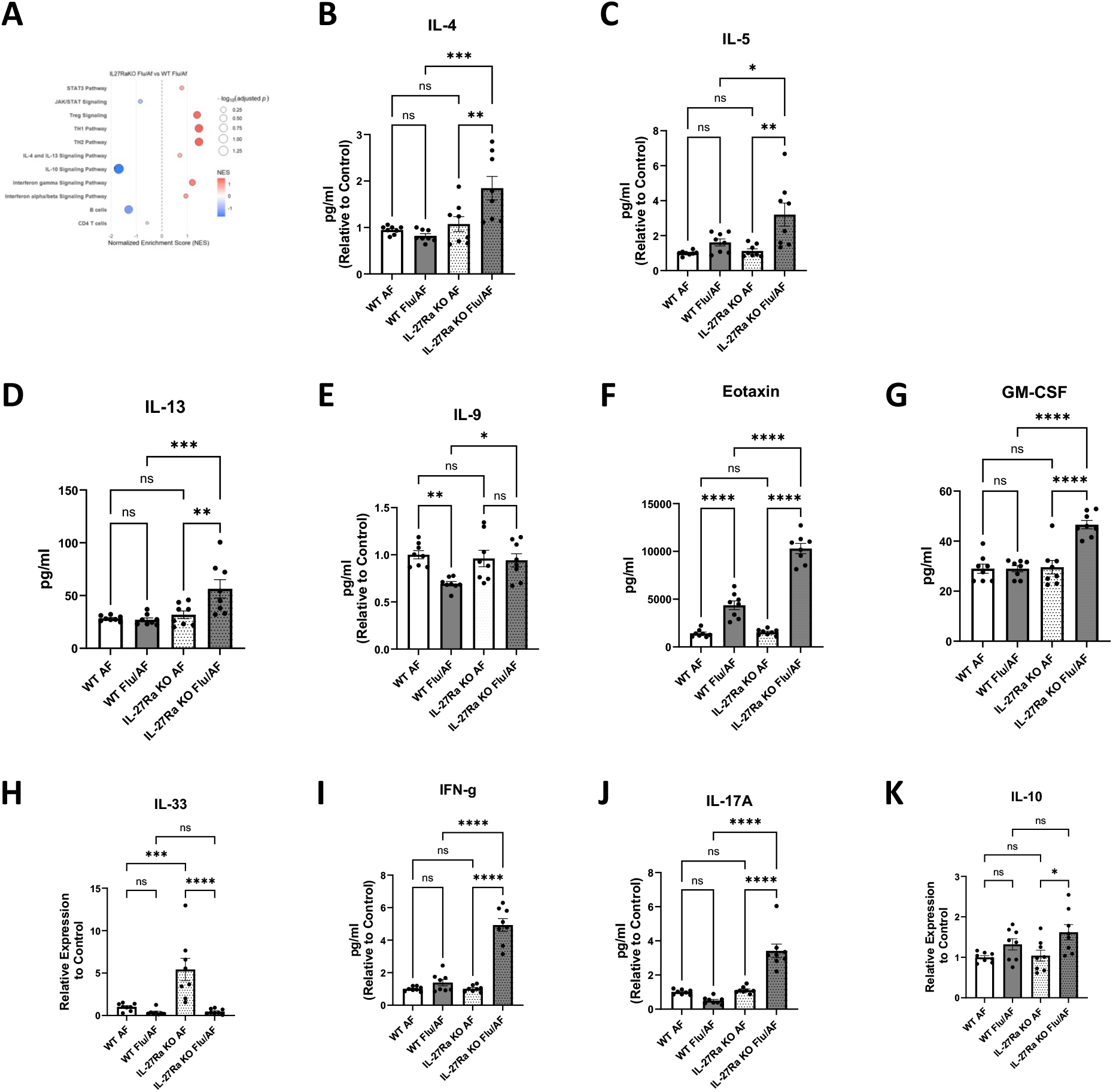
IL-27Rα deficiency enhances Type 2 immunity during IAPA. A-Gene Set Enrichment Analysis (GSEA) of bulk RNA sequencing data from IL-27Rα KO Flu/AF versus WT Flu/AF whole lung homogenate, using targeted immune signaling pathway gene sets from MSigDB (Mus musculus). Positive NES values indicate relative enrichment/upregulation in IL-27Rα KO Flu/AF compared with WT Flu/AF, whereas negative NES values indicate relative suppression/downregulation. Dot size corresponds to −log_10_(adjusted p-value) (n= 4 mice per group, data from one experiment).B-G-Lung protein levels of IL-4 (B), IL-5 (C), IL-13 (D), IL-9 (E), eotaxin (F), and GM-CSF (G) at 48 hours post-AF challenge, measured by multiplex immunoassay (Luminex). IL-4 (B), IL-5 (C), IL-9 (E), and eotaxin (F) are normalized to control (n= 8 mice per group).H-Relative *Il33* mRNA expression by qPCR in lungs normalized to control (n= 8 mice per group).I-J-Lung protein levels of IFN-γ (I) and IL-17A (J) measured by Luminex, normalized to control (n= 8 mice per group).K-Relative *Il10* mRNA expression by qPCR in lungs normalized to control (n= 8 mice per group) Data are shown as means ± SEM. *p<0.05, **p<0.005, ***p<0.0005, ****p<0.0001 by one-way ANOVA with Tukey’s multiple comparisons test (ns, not significant). All experiments represent a minimum of two independent experiments unless otherwise noted.

Next, we identified Type 2 immune cell types in the lung using flow cytometry and observed increased total gata3^+^ cells and Th2 cells (CD45 ^+^CD4^+^ GATA3^+^). Pathogenic Th2 cells, a highly inflammatory Th2 subset (CD4^+^ GATA3^+^ ST2^+^), and ILC2s (Lin^−^ GATA3^+^ ST2^+^) were also increased in IAPA *Il27ra^−/−^* mice compared to IAPA WT controls (Figure 4A-D). We quantified eosinophils, key downstream type 2 effector cell, and observed that both eosinophils (CD45^+^ Siglec-F^+^ CD11c^−/low^), and inflammatory eosinophils, a recruited, IL-5-dependent pro-inflammatory subset (Siglec-F^hi^ CD11c^lo^)^28^ are increased in IAPA *Il27ra^−/−^* mice compared to IAPA WT controls (Figure 4E-F). In parallel with cytokine levels and total cell numbers, transcriptomic profiling of the whole lung from IAPA *Il27ra^−/−^* mice revealed a marked shift toward type 2/eosinophil-associated programs. The chord diagram summarizes the differential expression of IAPA *Il27ra^−/−^* versus IAPA WT curated eosinophil modules, showing enrichment of eosinophil cell movement, eosinophil chemotaxis, and eosinophil activation pathways with associated genes displaying both increased and decreased expression, in the lungs of IAPA *Il27ra^−/−^* mice (Figure 4G). We hypothesized that antifungal effector functions may be altered and used the FLARE assay to quantify conidial uptake and viability of conidia within eosinophils(23). Lung eosinophils and inflammatory eosinophils from IAPA *Il27ra^−/−^*mice both exhibited reduced killing relative to IAPA WT controls, indicating that despite their increased abundance, these cells are functionally impaired in conidial killing (Figure 4 H-I). We did not observe any differences in conidial uptake by eosinophils and inflammatory eosinophils (Figure 4J-K). When eosinophil subsets were directly compared, inflammatory eosinophils (Siglec-F^hi^ CD11c^lo^) showed decreased conidial uptake but increased killing of *A. fumigatus* conidia compared to eosinophils (CD45^+^ Siglec-F^+^ CD11c^−/low^) in IAPA WT mice (Figure 4L-M). These data suggest that IL-27Rα deficiency leads to enhanced numbers of eosinophils that have reduced functional capacity to kill *A. fumigatus* during IAPA, which may contribute to the increased fungal burden observed in these mice. Additionally, the subset of inflammatory eosinophils is functionally altered compared to eosinophils with increased conidial killing but decreased conidial uptake of conidia.

**Figure 4.**
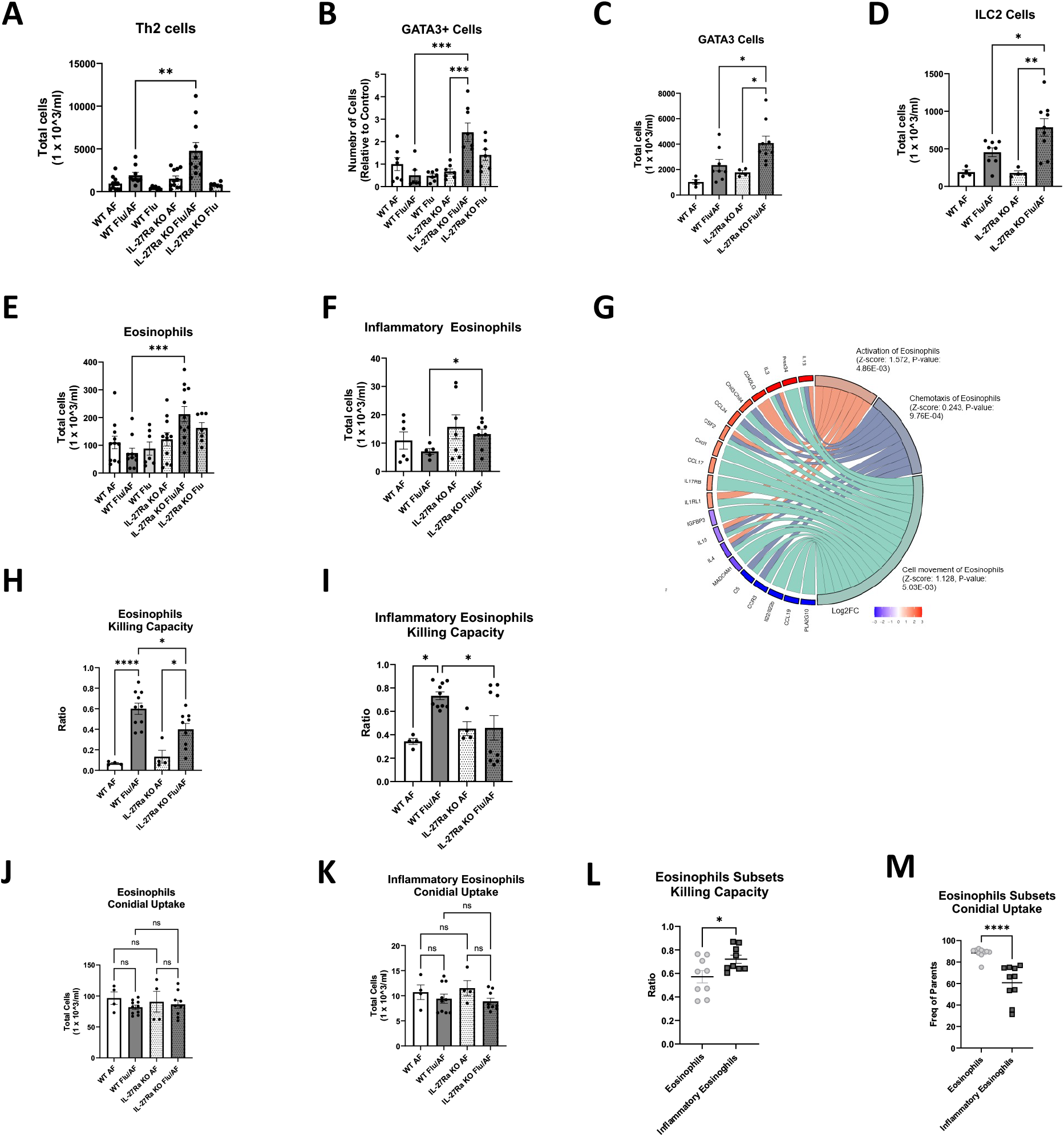
IL-27Rα deficiency expands type 2 lymphocytes and eosinophils with impaired antifungal killing during IAPA. A-D-Total number of Th2 cells (CD45^+^CD4^+^GATA3^+^) (A), GATA3^+^ cells (B), Pathogenic Th2 cells (CD4^+^GATA3^+^ST2^+^) (C), and ILC2s (Lin^−^GATA3^+^ST2^+^) (D) in lungs of WT AF, WT Flu/AF, WT Flu, IL-27Rα KO AF, IL-27Rα KO Flu/AF, and IL-27Rα KO Flu groups by flow cytometry (n= 6-12 mice per group). E-F-Total eosinophils (CD45^+^Siglec-F^+^CD11c^−^/lo) (E) and inflammatory eosinophils (Siglec-F^hi CD11c^lo) (F) in lungs by flow cytometry (n= 7-12 mice per group).G-Chord diagram of IPA (Qiagen)-curated eosinophil pathway analysis comparing IL-27Rα KO Flu/AF versus WT Flu/AF lungs. Pathways shown include eosinophil activation (Z-score: 1.572, p=4.86×10^−3^), chemotaxis of eosinophils (Z-score: 0.243, p=9.76×10^−4^), and cell movement of eosinophils (Z-score: 1.128, p=5.03×10^−3^). Positive Z-scores indicate activation/upregulation in IL-27Rα KO Flu/AF compared with WT Flu/AF; negative Z-scores indicate inhibition/downregulation. Chords connect pathways to their associated differentially expressed genes; gene color indicates Log2FC in IL-27Rα KO Flu/AF compared with WT Flu/AF (n= 4 mice per group, data from one experiment).H-I-Conidial killing capacity of total eosinophils (H) and inflammatory eosinophils (I) in WT AF, WT Flu/AF, IL-27Rα KO AF, and IL-27Rα KO Flu/AF mice, assessed by FLARE assay (n= 4-10 mice per group).J-K - Conidial uptake of total eosinophils (J) and inflammatory eosinophils (K) in WT AF, WT Flu/AF, IL-27Rα KO AF, and IL-27Rα KO Flu/AF mice, assessed by FLARE assay (n= 4-10 mice per group). L-M-Conidial killing capacity (L) and conidial uptake (M) comparing total eosinophils versus inflammatory eosinophils within IAPA WT mice (n= 9 mice per group). Data are shown as means ± SEM. *p<0.05, **p<0.005, ***p<0.0005, ****p<0.0001 by one-way ANOVA with Tukey’s multiple comparisons test (A–F, H–K) and unpaired t-test (L–M) (ns, not significant). All experiments represent a minimum of two independent experiments unless otherwise noted.

### IL-27Rα deficiency decreases macrophage numbers and limits macrophage-mediated conidial uptake is during IAPA

As macrophages are key pulmonary effector cells against *A. fumigatus*(26)(27), we hypothesized that macrophage numbers and function may be altered by the deletion of the IL-27Rα in mice. Lung RNA-sequencing pathway analysis (*Il27ra^−/−^* IAPA vs WT IAPA) demonstrated inhibition of Fcγ receptor–macrophage mediated phagocytosis, quantity of macrophages, and macrophage classic activation pathways, while macrophage alternative activation pathway, cell movement of macrophages, production of NO and ROS pathways were upregulated (Figure 5A). To quantify macrophages in bronchoalveolar lavage fluid, we performed cyto-spin differential cell counting and observed decreased total numbers of macrophages in the IAPA *Il27ra^−/−^* mice compared to IAPA WT controls (Figure 2 F). Next, we quantified macrophages in the lung by flow cytometry and observed decreased total macrophage numbers in IAPA *Il27ra^−/−^* mice compared to IAPA WT controls (Figure 5B). To assess macrophage function, we used FLARE conidia to quantify macrophage conidial uptake and killing capacity and observed decreased conidial uptake of conidia by macrophages, although there was no change in killing capacity of conidia by macrophages (Figure 5C,D). Next, we examined macrophages subsets the lung, alveolar macrophages and interstitial macrophages, both of which were decreased in abundance in *Il27ra^−/−^* IAPA mice compared to WT control (Figure 5E, H). Killing capacity and conidial uptake of alveolar macrophages were unchanged between *Il27ra^−/−^* and WT IAPA mice (Figure 5F, G) while interstitial macrophages showed no change in killing capacity but a significant decrease in conidial uptake (Figure 5I, J). We also examined macrophages based on activation state, defined as classical activation (M1), alternative activation (M2), or mixed M1/M2(28). Although we hypothesized that there would be increased M2 macrophages in the *Il27ra^−/−^* IAPA mice based on cytokine and lung RNA seq analysis, we observed no differences in total numbers or function of these different types of macrophages (Figure 5K-M). Collectively, these data suggest during IAPA that IL-27Rα deficiency primarily affects macrophage uptake rather than intrinsic killing and reduced macrophage abundance and impaired conidial uptake capacity in *Il27ra^−/−^* IAPA mice may contribute to the observed increased fungal burden. These data support IL-27Rα-dependent regulation of macrophage antifungal responses during IAPA.

**Figure 5.**
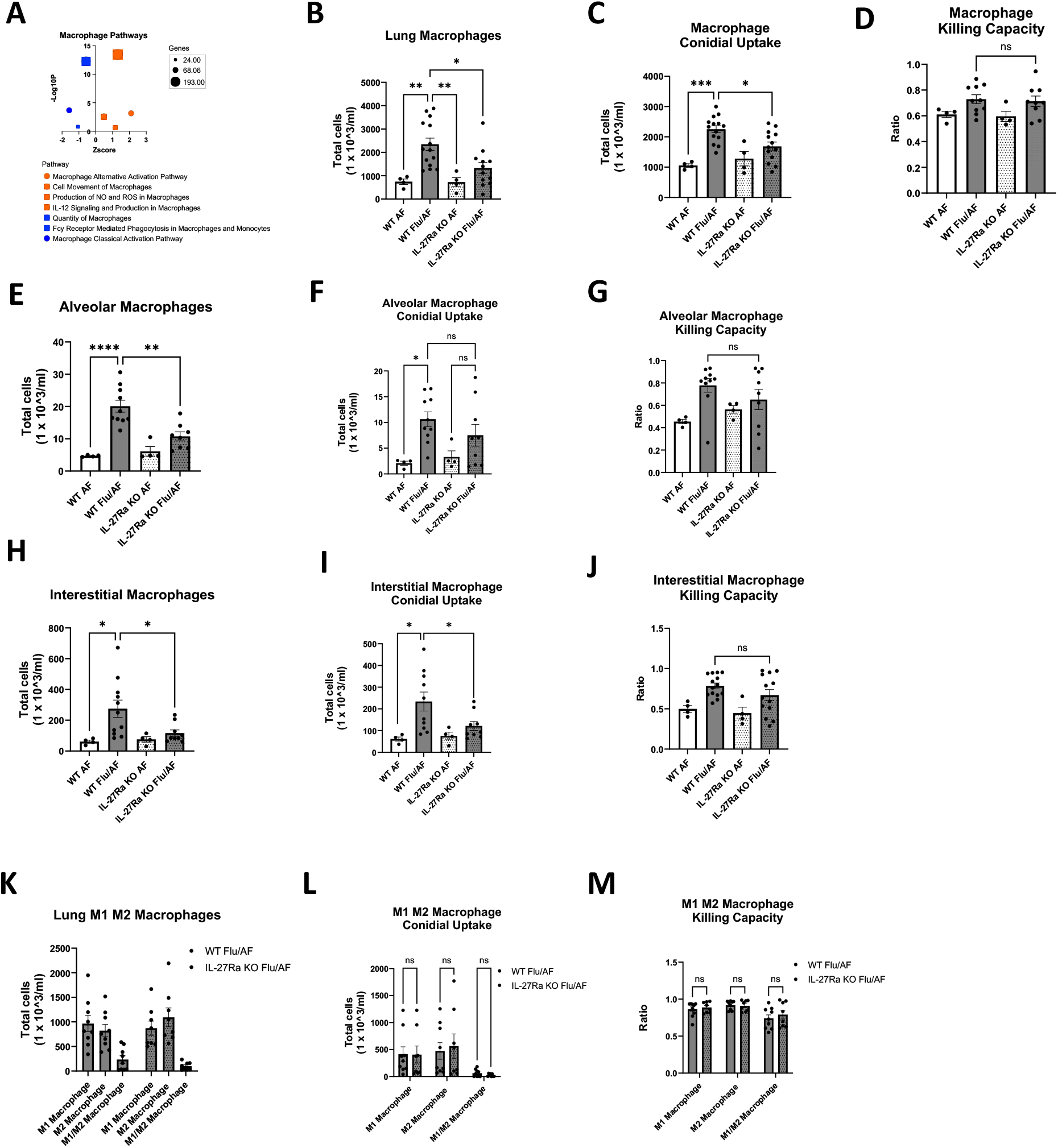
IL-27Rα deficiency decreases macrophage numbers and limits macrophage-mediated conidial uptake during IAPA. A-Dot plot of macrophage-related pathway analysis from IPA (Qiagen) comparing IL-27Rα KO Flu/AF versus WT Flu/AF lungs. Positive Z-scores indicate activation/upregulation and negative Z-scores indicate inhibition/downregulation in IL-27Rα KO Flu/AF compared with WT Flu/AF. Dot size corresponds to −log_10_(p-value) (n= 4 mice per group, data from one experiment).B-D-Total lung macrophage counts (B), conidial uptake (C), and killing capacity (D) (n= 4-14 mice per group). E-G-Alveolar macrophage counts (E), conidial uptake (G), and killing capacity (I) (n= 4-14 mice per group). H-J-Interstitial macrophage counts (H), conidial uptake (I), and killing capacity (J) (n= 4-14 mice per group). K-M-Total numbers of M1, M2, and mixed M1/M2 macrophage subsets (K), conidial uptake (L), and killing capacity (M) (n= 9 mice per group). Data are shown as means ± SEM. *p<0.05, **p<0.005, ***p<0.0005, ****p<0.0001 by one-way ANOVA with Tukey’s multiple comparisons test (B–L) and unpaired t-test (A, M–O) (ns, not significant). All experiments represent a minimum of two independent experiments unless otherwise noted.

### Exogenous IL-27 improves fungal clearance and survival during IAPA

To determine whether augmentation of IL-27 signaling alters antifungal host defense during IAPA, WT mice received intraperitoneal rIL-27 on days 4-7 of our IAPA model, whereas control mice received PBS sham (Figure 6A). rIL-27 treatment enhanced fungal clearance during IAPA, demonstrated by both reduced CFU in lung tissue compared with PBS-treated IAPA controls and quantification of fungal burden on GMS-stained lung sections (Figure 6 B-D). Moreover, rIL-27 treatment significantly reduced influenza viral burden compared to PBS-treated WT Flu/AF controls (Figure 6E). rIL-27 significantly improved survival compared with PBS-sham IAPA controls (Figure 6F). Consistent with the established role of IL-27 in inhibition of type 2 immunity, rIL-27 treatment also modulated eosinophil responses during IAPA. Total lung eosinophils (CD45^+^ Siglec-F^+^ CD11c^−/low^) and total lung inflammatory eosinophils (Siglec-F^hi^ CD11c^lo^)^28^ were reduced. The killing capacity of eosinophils was increased in mice that received rIL-27, although there were no changes in conidial uptake by eosinophils (Figure 6G-J). Together, these findings indicate that improved outcomes with rIL-27 are accompanied by enhanced fungal clearance and altered type 2–associated effector responses.

**Figure 6.**
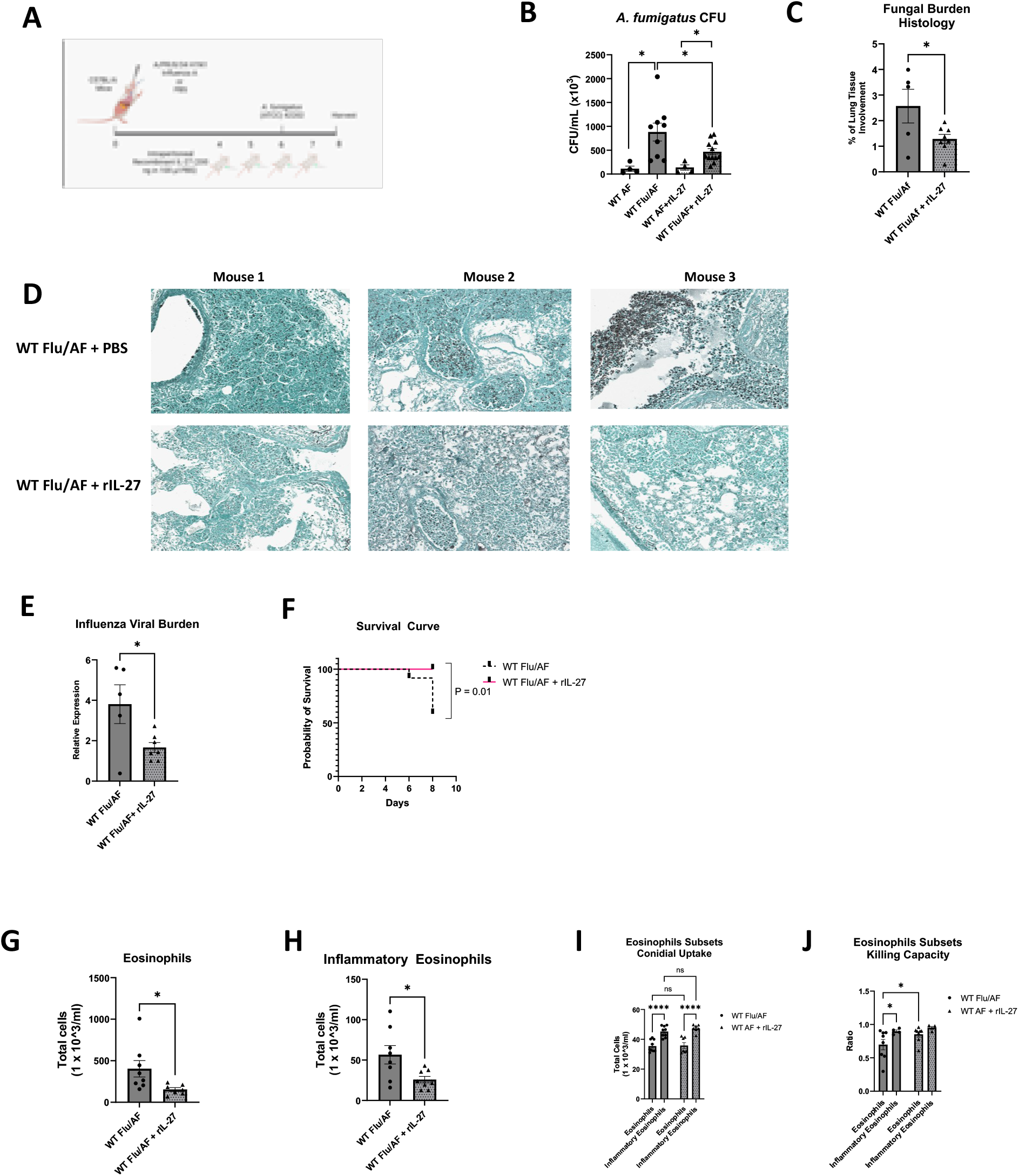
Timed recombinant IL-27 administration improves fungal clearance and survival during IAPA. A-Schematic of the IAPA mouse model and rIL-27 treatment protocol. C57BL/6 mice were infected with influenza A/PR/8/34 H1N1 or PBS on day 0, challenged with *A. fumigatus* (ATCC 42202) on day 6, and received intraperitoneal recombinant IL-27 (200 ng in 100 µL PBS) or PBS on days 4–7, with lung harvest on day 8. B-*A. fumigatus* CFU/mL (×10^3^) in lung homogenates (n= 4-12 mice per group). C-Digital quantification of pulmonary *A. fumigatus* burden on GMS-stained lung sections, expressed as percentage of lung tissue area involvement (n= 5-7 mice per group). D-Representative GMS-stained lung sections from three mice per group, demonstrating reduced fungal burden with rIL-27 treatment. E-Relative influenza A/PR/8/34 viral burden by qPCR in lungs (n= 5-7 mice per group). F-Survival curve of WT Flu/AF and WT Flu/AF+rIL-27 mice over 8 days (log-rank Mantel-Cox test, p=0.01) (n= 12 mice per group). G-HTotal eosinophils (CD45^+^Siglec-F^+^CD11c^−^/lo) (G) and inflammatory eosinophils (Siglec-F^hi CD11c^lo) (H) in lungs (n= 8 mice per group). I-J-Conidial uptake (I) and Conidial killing capacity (J) of total eosinophils and inflammatory eosinophils compared side by side (n= 8 mice per group). Data are shown as means ± SEM. *p<0.05, **p<0.005, ***p<0.0005, ****p<0.0001 by unpaired t-test (B, C, E, G–L) and log-rank Mantel-Cox test (E) (ns, not significant). All experiments represent a minimum of two independent experiments unless otherwise noted.

### Exogenous IL-27 promotes alveolar macrophage conidial uptake during IAPA

Next, we investigated if enhancement of IL-27 signaling alters macrophage abundance and antifungal function during IAPA. In mice that received rIL-27 on Days 4-7 of our IAPA model, we observed no differences in total numbers of lung macrophages, alveolar macrophages, or interstitial macrophages (Figure 7A, D, G). Using FLARE conidia to quantify uptake and killing, we found that conidial uptake was increased in mice that received rIL-27 during IAPA compared to those that received a PBS sham (Figure 7B). When we assessed the conidial uptake within macrophages located in different anatomical compartments, we observed increased conidial uptake by alveolar macrophages following timed rIL-27 administration, whereas there was no change in conidial uptake by interstitial macrophages (Figure 7E, H). We observed no differences in fungal killing by alveolar or interstitial macrophages (Figure 7C, F, I). We also examined macrophages based on activation state: M1, M2, or mixed M1/M2, and observed an increase of total numbers of mixed M1/M2 macrophages but no differences in killing capacity or conidial uptake by these macrophages (Figure 7J-L).

**Figure 7.**
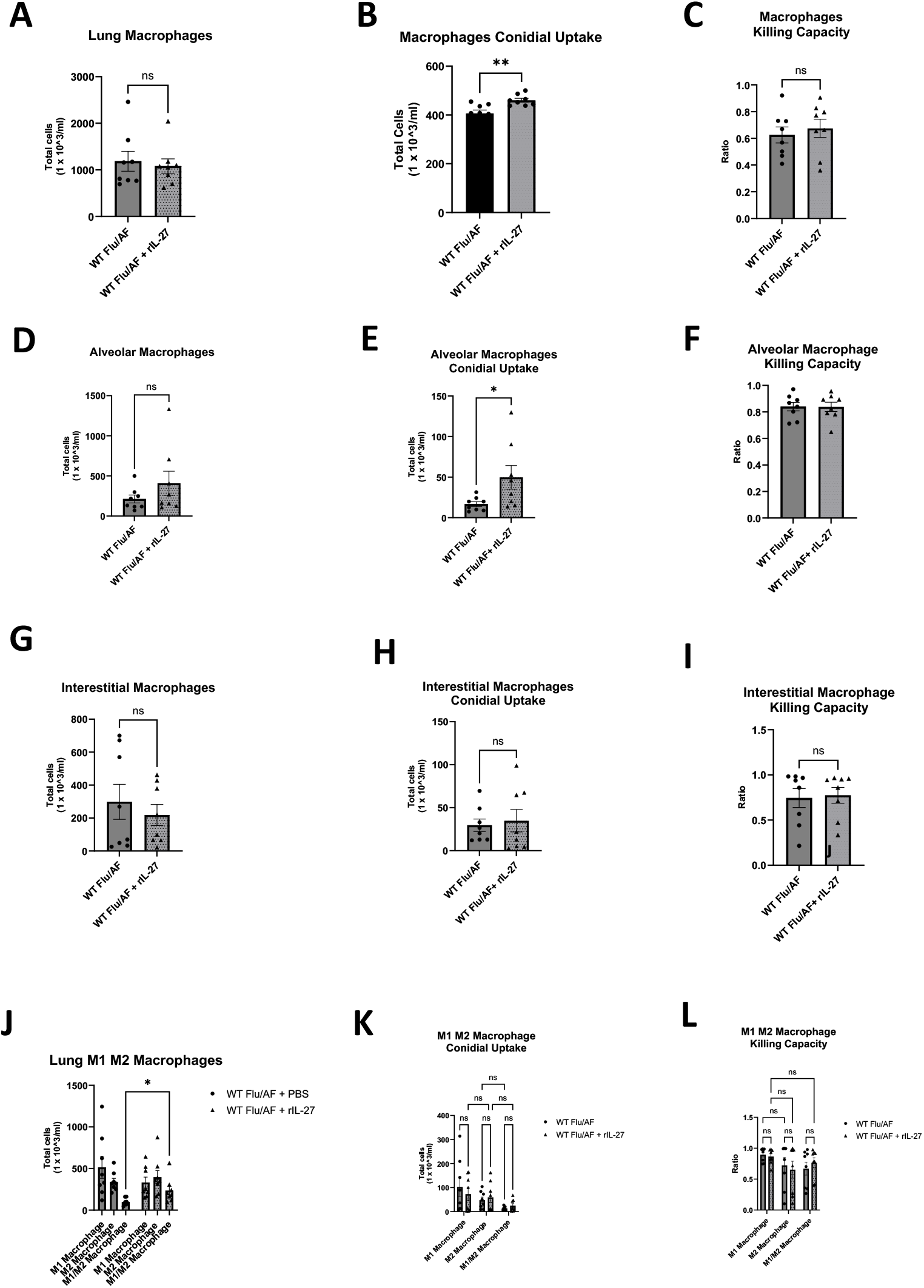
Recombinant IL-27 promotes macrophage-mediated conidial uptake during IAPA. A-C-Total lung macrophage counts (A), conidial uptake (B), and killing capacity (C) (n= 8 mice per group). D-F-Alveolar macrophage counts (D), conidial uptake (E), and killing capacity (F) (n= 8 mice per group).G-I-Interstitial macrophage counts (G), conidial uptake (H), and killing capacity (I) (n= 8 mice per group).J-L-Total numbers of M1, M2, and mixed M1/M2 macrophage subsets (J), conidial uptake(n= 8 mice per group). Data are shown as means ± SEM. *p<0.05, **p<0.005, ***p<0.0005, ****p<0.0001 by unpaired t-test (A–I) and one-way ANOVA with Tukey’s multiple comparisons test (J–L) (ns, not significant). All experiments represent a minimum of two independent experiments unless otherwise noted.

## Discussion

In this study, we identify IL-27 signaling as a protective cytokine during IAPA. *Il27ra^−/−^* IAPA mice demonstrated significantly higher pulmonary fungal burden compared to WT IAPA mice. This susceptibility phenotype was functionally supported by gain-of-function studies, as timed administration of rIL-27 enhanced fungal clearance and improved survival during IAPA. After establishing this protective role of IL-27 signaling, we sought to define downstream effector mechanisms, focusing on aspects of the immune system known to be affected by IL-27.

IL-27 signaling through IL-27Rα shapes pulmonary antifungal host defense in a pathogen- and context-dependent manner. In systemic infection with *Candida parapsilosis*, loss of IL-27R signaling is associated with enhanced fungal clearance, whereas in pulmonary cryptococcosis due to *Cryptococcus neoformans*, deficiency of IL-27 signaling is reported to increase lung fungal burden and shorten survival ^32,33^. During pulmonary aspergillosis, published studies likewise report divergent roles for IL-27 signaling across different experimental settings. Strickland and colleagues used a repeated-exposure model in immunocompetent mice, delivering *A. fumigatus* 2.5 × 10^7^ Af293 conidia intranasally every other day for 28 days, and found that IL-27Rα deficiency increased lung fungal burden, supporting a protective role for IL-27 signaling in that setting^34^. In contrast, Liu and colleagues employed an acute invasive pulmonary aspergillosis model using a single intratracheal challenge with *A. fumigatus* Af293 and reported that IL-27Rα deficiency developed less severe infection with decreased fungal burden, reduced lung pathology, and improved survival^19^. In our current experiments using a different *A. fumigatus* strain, we observed comparable results to Liu et al between *Il27ra^−/−^* and WT mice in the single acute *A. fumigatus* infection setting, suggesting that the impact of IL-27 signaling likely differs between chronic and acute exposure to *A. fumigatus* in the lung. However, acute infection with IAPA revealed a different phenotype, as IL-27Rα deficiency was associated with significantly increased pulmonary fungal burden compared with WT IAPA mice. During influenza infection, IL-27 protects from immunopathology, with *Il27ra^−/−^* exhibiting an altered immune response but no differences in viral burden ^35,36^. Notably, we observed increased influenza burden in *Il27ra^−/−^* IAPA mice compared to WT mice. Interestingly, IL-27 signaling has been studied in models of post-influenza bacterial super-infection, including *Streptococcus pneumoniae* and *Staphylococcus aureus* ^35,37^, and decreased bacterial burden was observed in *Il27ra^−/−^* mice compared to WT controls, showing that IL-27 signaling plays disparate roles in post-influenza fungal and bacterial secondary pneumonia.

Mechanistically, IL-27 signaling activates a STAT1/STAT3-dependent transcriptional axis that favors Type 1 polarization, including the induction of T-bet and promotion of Th1 differentiation, while suppressing Th2 and Th17 development ^38-40^. IL-27 suppresses Th2 differentiation and Th2 cytokine production, in part through STAT1-dependent downregulation of GATA-3 and can inhibit cytokine output from polarized Th2 cells ^16,38^. In addition, IL-27 directly restrains innate type 2 responses by inhibiting ILC2 proliferation and IL-5/IL-13 production, thereby limiting downstream eosinophilic airway inflammation ^17,41,42^. In our current study, in *Il27ra^−/−^* IAPA mice, Type 2 immunity was broadly amplified, with increased Th2 cells, pathogenic Th2 cells, and ILC2 cells compared to WT IAPA mice. This cellular expansion was mirrored by a coordinated Type 2 cytokine signature, with IL-4, IL-5, IL-9, and IL-13 increased in lungs of *Il27ra^−/−^* IAPA mice. Interestingly, IL-33 was not different between *Il27ra^−/−^* and WT IAPA mice, although there was a strong induction of IL-33 during *A. fumigatus* infection in *Il27ra^−/−^* mice compared to WT, something that can be explored further in future studies. Prior studies have shown that IL-4 deficient mice were more resistant to pulmonary *A. fumigatus* infection(29), however, IL-33 deficient mice have enhanced clearance of *A. fumigatus* from the lung(30), suggesting that individual Type 2 cytokines have different roles during invasive pulmonary aspergillosis. Additionally, IL-4Rα and STAT6 partially regulate arginase gene expression, the prototypic marker for alternatively activated macrophages, in alveolar macrophages during acute pulmonary *A. fumigatus* infection(31).

Prior literature demonstrates a direct contribution of eosinophils to antifungal host defense. Eosinophil deficiency compromises pulmonary clearance of *A. fumigatus*, indicating that eosinophils can participate in fungal control in vivo^19^. Using FLARE conidia, Guerra and colleagues showed that lung eosinophils associate with conidia and mediate measurable conidial killing *in vivo* ^43^. In our current studies, downstream effector Type 2 immune cells, total eosinophils, and inflammatory eosinophils were increased in *Il27ra^−/−^* IAPA mice compared to WT IAPA mice. Although there was an increased abundance of eosinophils in *Il27ra^−/−^* IAPA mice, the killing capacity of the total eosinophil population was reduced in *Il27ra^−/−^* IAPA mice compared to WT IAPA mice. Notably, inflammatory eosinophils had reduced conidial uptake and increased killing capacity compared to conventional eosinophils, a finding that expands upon the current knowledge regarding the functional capacity of inflammatory eosinophils and can be explored further in future studies. Timed administration of rIL-27 reduced eosinophil abundance during IAPA but enhanced fungal killing. Together, these findings support the interpretation that loss of IL-27 signaling allows for an exaggerated type 2 polarization with increased eosinophilia during IAPA; however, eosinophil effector function and killing capacity of conidia is reduced, potentially resulting in increased fungal burden on the. IL-5 and GM-CSF are primary eosinophil-activating cytokines that are predicted to enhance eosinophil antifungal killing; however, there is limited data linking any specific cytokines to eosinophil intracellular killing of *A. fumigatus* conidia. Both IL-5 and GM-CSF are increased in *Il27ra^−/−^* IAPA mice compared to WT IAPA mice and future studies will need to address the mechanism by which eosinophils have reduced fungal killing of aspergillus conidia in IL-27Rα deficient mice.

Although studies using FLARE conidia in IAPA mouse models have not revealed differences in conidial uptake or killing by wild-type IAPA macrophages compared to singular *A. fumigatus* infection ^5,44^, our current experiments reveal a IL-27Rα–signaling dependent effect on macrophage antifungal effector function. IAPA in *Il27ra^−/−^* mice exhibited reduced alveolar and interstitial macrophage abundance and decreased conidial uptake of conidia by interstitial macrophages compared to WT IAPA mice. Conversely, therapeutic administration of rIL-27 enhanced conidial uptake by alveolar macrophages during IAPA. Surprisingly, we observed minimal differences in M1, M2, or mixed M1/M2 macrophage abundance and function during IAPA. Together, these findings support the interpretation that IL-27 signaling enhances macrophage effector function during IAPA. A notable limitation of the use of FLARE conidia is that they are most well suited to examine early innate immune responses before conidial germination and hyphal formation.

Prior studies during *A. fumigatus* infection have established a relationship between eosinophils and macrophages. Eosinopenic mice exhibit reduced recruitment of inflammatory monocytes and diminished expansion of lung macrophage populations after *A. fumigatus* conidia challenge, suggesting eosinophils can also influence broader myeloid organization during acute pulmonary aspergillosis^43^. Future studies will need to be performed to determine cell-specific contributions of eosinophils and macrophages to IAPA pathogenesis and to determine if there is a dependent or independent relationship between eosinophils and macrophages during IAPA. Using single cell sequencing, we have demonstrated that T cells are the major recipients of IL-27 signaling and observed higher expression of *Il27ra* in IAPA T cells compared to singular *A. fumigatus.* Differential expression of the IL-27 receptor on T cells likely leads to downstream effector cytokines that impact eosinophil and macrophage function. A potential alternative interpretation is that an autocrine look exists within monocytes. Additional work is needed to determine which specific cytokines regulate eosinophil and macrophage function during IAPA and Type 2 cytokines may not be the only driving force behind protective IL-27 signaling during IAPA.

In summary, IL-27 signaling is protective during IAPA. Timed administration of exogenous IL-27 near the peak of viral load during singular influenza infection protects against an exaggerated inflammatory response (32) and timed administration of exogenous IL-27 near the peak of viral load during IAPA improves survival, reduces fungal and viral burden, and enhances eosinophil killing and macrophage uptake of fungal conidia. Currently, human anti-IL-27 antibodies are being studied in clinical trials for solid tumors (33), and it has also been proposed that human recombinant IL-27 may be therapeutic for its anti-tumor activity by T cells(34). T cells have been engineered to express IL-27 and shown anti-tumor activity with decreased pro-inflammatory cytokines (35). Infectious complications could be increased with the use of human anti-IL-27 antibodies; however, the benefit of tumor reduction will likely outweigh potential risks and may carry less risk infectious than other cancer treatments. Future studies will need to address the timing, dosage, and side effects of recombinant IL-27 as a therapeutic agent during IAPA.

## Author contributions

NN assisted in study design [supporting], performed experiments [lead], analyzed data [equal], wrote the manuscript [equal]. YKM performed experiments [supporting], analyzed data [supporting], and assisted with editing the manuscript [supporting]. SW, RBS, and CN performed experiments [supporting]. DL [equal] and KC [equal] performed the scRNA-seq analyses. RG provided the influenza virus [lead] and assisted with editing the manuscript [supporting]. KMR conceptualized and designed the studies [lead], analyzed data [equal], wrote the manuscript [equal], and supervised the studies [lead]. All authors reviewed the manuscript.

## Acknowledgment

FLARE conidia were kindly provided by Dr. Tobias Hohl (Memorial Sloan Kettering Cancer Center). This project utilized the services of the University of Pittsburgh Single Cell Core Facility (RRID:SCR_025110), supported in part by the University of Pittsburgh Office of the Senior Vice Chancellor for Health Sciences. This project utilized the services of the University of Pittsburgh Health Sciences Sequencing Core at UPMC Children’s Hospital of Pittsburgh, for bulk RNA sequencing.

## Funding

Funding sources include NIAID R01AI153337 (KMR), NHLBI R03HL154242 (KMR), and NHLBI R01 HL146479 (RG)

## Conflicts of Interest

None Declared

## Data availability

The single-cell RNA-seq data will be deposited into the GEO database at the time of publication acceptance. All other raw data are available from the lead contact upon request.

## Notes

### Competing Interest Statement

The authors have declared no competing interest.

