## Supplemental Figures and Tables for "IL-27 Signaling Protects Against Influenza-Associated Pulmonary Aspergillosis Through Inhibition of Type 2 Immunity and Enhanced Antifungal Immunity"

Figure S1

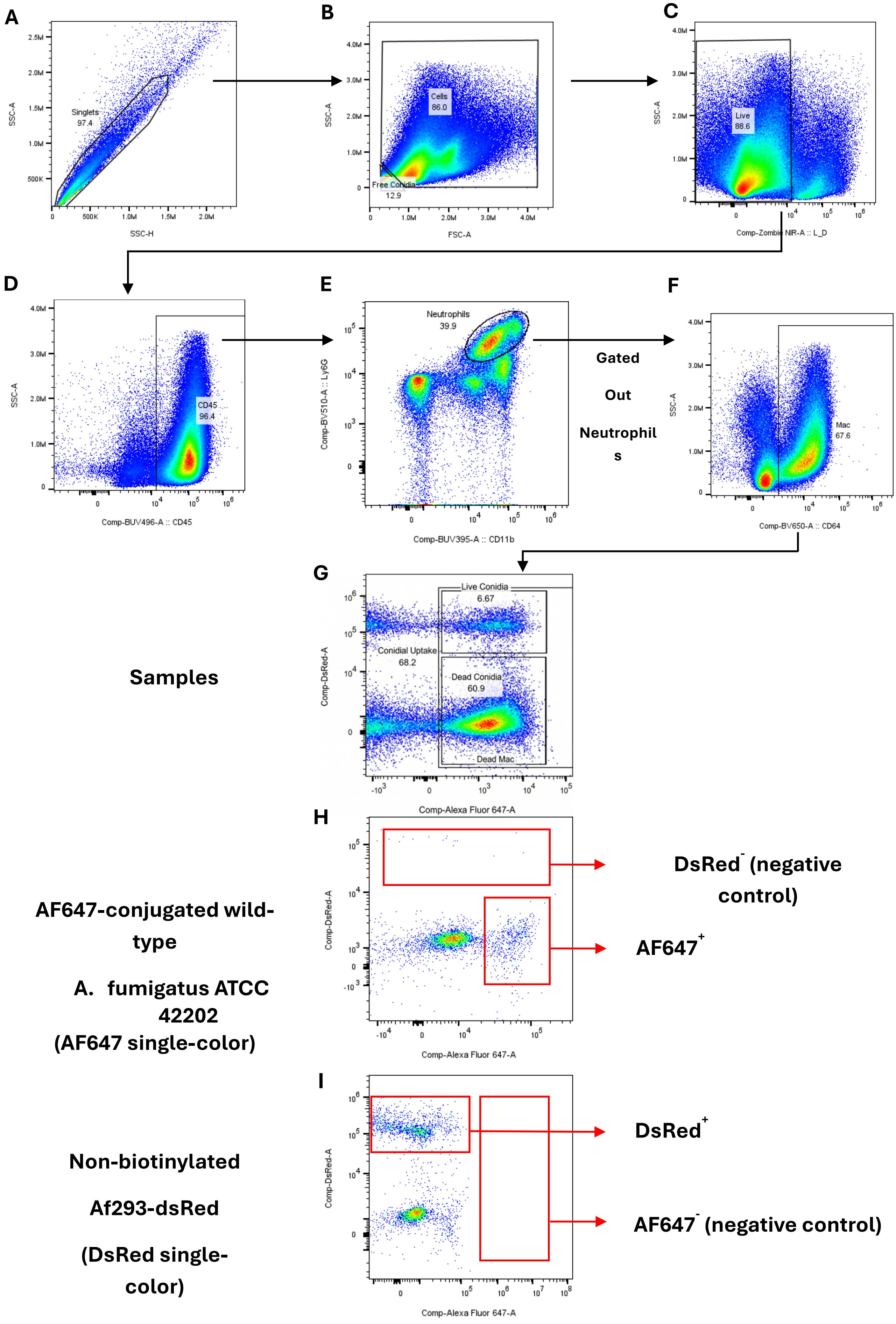
A
Singlets 97.4
SSC-A
SSC-H
B
Cells 86.0
Free Conidia 12.9
SSC-A
FSC-A
C
Live 88.6
SSC-A
Comp-Zombie NIR-A :: L_D
D
CD45 96.4
SSC-A
Comp-BUV496-A :: CD45
E
Neutrophils 39.9
Comp-BV510-A :: Ly6G
Comp-BUV395-A :: CD11b
Gated Out Neutrophils
F
Mac 67.6
SSC-A
Comp-BV650-A :: CD64
G
Samples
Live Conidia 6.67
Conidial Uptake 68.2
Dead Conidia 60.9
Dead Mac
Comp-DsRed-A
Comp-Alexa Fluor 647-A
H
AF647-conjugated wild-type
A. fumigatus ATCC 42202
(AF647 single-color)
DsRed- (negative control)
AF647+
Comp-DsRed-A
Comp-Alexa Fluor 647-A
I
Non-biotinylated
Af293-dsRed
(DsRed single-color)
DsRed+
AF647- (negative control)
Comp-DsRed-A
Comp-Alexa Fluor 647-A

### Figure S1 – Flow Cytometry Gating Strategy

Lung cells were analyzed 48 hours following *A. fumigatus* FLARE conidia. Lung single-cell suspensions were prepared as described in the Methods. **(A)** Singlets were identified by SSC-H vs SSC-A gating. **(B)** Free *Aspergillus fumigatus* conidia were gated as $FSC^{lo}$ $SSC^{lo}$ events and excluded; immune cells were selected from the remaining population. **(C)** Live/dead discrimination was performed using Zombie NIR. **(D)** $CD45^{+}$ leukocytes were gated. **(E)** Neutrophils were identified as $CD45^{+}$ $Ly6G^{+}$ $CD11b^{+}$ cells. **(F)** Neutrophils were excluded, and macrophages were subsequently identified within the $CD45^{+}$ $Ly6G^{-}$ population as $CD64^{+}$ cells. **(G)** Conidial uptake was determined by AF647 positivity. $DsRed^{+}$ $AF647^{+}$ cells represent macrophages containing live conidia, whereas $DsRed^{-}$ $AF647^{+}$ cells indicate macrophages containing dead conidia. Each flowcytometry experimental run included two compensation controls: a mouse challenged with AF647-conjugated wild-type A. fumigatus ATCC 42202 to serve as the AF647 single-color reference (H), and a mouse challenged with non-biotinylated Af293-dsRed conidia to serve as the DsRed single-color reference (I). These controls were processed in parallel with experimental samples to ensure accurate spectral unmixing and population gating.

**Table S1**

| Target | Fluorophore | Clone | Vendor |
|---|---|---|---|
| Lineage cocktail (CD3/Ly6G-Ly6C/CD11b/CD45R(B220)/TER-119) | Alexa Fluor 700 | 17A2/RB6-8C5/M1/70/RA3-6B2/TER-119 | BioLegend |
| Ly6G | BV510 | 1A8 | BioLegend |
| F4/80 | BV480 | T45-2342 | BD Horizon |
| Siglec F (CD170) | PerCP-eFluor 710 | 1RNM44N | Invitrogen |
| CD11b | BUV395 | M1/70 | BD Horizon |
| CD45 (1) | Alexa Fluor 532 | 30-F11 | Invitrogen |
| CD64 | BV650 | X54-5/7.1 | BioLegend |
| CD45 (2) | BUV496 | 30-F11 | BD OptiBuild |
| CD4 | Pacific Blue | RM4-5 | BD Pharmingen |
| GATA3 | Alexa Fluor 488 | 16E10A23 | BioLegend |
| CD86 | BUV737 | GL1 | Invitrogen |
| CD206 | BV605 | C068C2 | BioLegend |
| ST2 (IL-33R) | PerCP-eFluor 710 | RMST2-2 | eBioscience |
| CD11c | PE-Fire 640 | N418 | BioLegend |

**Table S1– Antibody panel**

Antibodies used for flow cytometric analysis of lung leukocyte populations, listed with target antigen, conjugated fluorophore, clone, and vendor. Two CD45 conjugates (Alexa Fluor 532 and BUV496) were used across panels, with the choice depending on which channel was otherwise occupied by another marker in that panel; panels were designed to avoid spectral overlap.
